# Integrated Proteomic and Epigenomic Analysis Reveals IGF2 as a Vulnerability in PRC2-Deficient Malignant Peripheral Nerve Sheath Tumors

**DOI:** 10.1101/2025.11.05.686792

**Authors:** Joanna K. Lempiäinen, Kirill Miachin, Xingyu Liu, Kuangying Yang, Cynthia Horth, Eric Bareke, Mitchell F. Grinwald, Wesley N. Saintilnord, Ting Wang, Jacek Majewski, Angela C. Hirbe, Benjamin A. Garcia

## Abstract

Malignant peripheral nerve sheath tumors (MPNSTs) are aggressive sarcomas with limited therapeutic options. Loss of the Polycomb repressive complex 2 (PRC2), via inactivating mutations in SUZ12 or EED, occurs frequently in MPNSTs and is associated with poor prognosis. However, the downstream chromatin and signaling consequences of these mutations remain incompletely understood. Here, we show that PRC2 deficiency in MPNST cells induces coordinated chromatin remodeling, characterized by loss of repressive H3K27me3 and gain of activating marks, including H3K27ac and H3K36me2. Integrative epigenomic, transcriptomic, and proteomic profiling revealed that this chromatin reprogramming activates a fetal-like growth signature centered on insulin-like growth factor 2 (IGF2) and its post-transcriptional regulators, Insulin-like Growth Factor 2 mRNA-Binding Protein (IGF2BP1-3). Functional studies demonstrate that PRC2-deficient cells are selectively dependent on IGF2 for proliferation, and that restoration of SUZ12 suppresses IGF2 expression and reduces growth. Analysis of human MPNST tumors confirms upregulation of the IGF2-IGF2BP axis in PRC2-deficient tumors, highlighting its clinical relevance. Together, these findings link PRC2 loss to activation of fetal growth factor-driven oncogenic signaling and identify IGF2 and its regulatory network as potential vulnerabilities in this aggressive tumor subtype.

## Introduction

Malignant peripheral nerve sheath tumors (MPNSTs) are clinically aggressive, high-grade soft tissue sarcomas with recurrence rates of approximately 50% and overall survival of less than five years despite surgery, radiation, and chemotherapy^1–3^. Composed of neoplastic Schwann cells, MPNSTs account for roughly 5% of all soft tissue sarcomas. The only curative approach remains aggressive local control through surgery and radiation therapy. Advanced disease typically follows a rapidly progressive course, with frequent metastases to the lungs and bone^4^. For patients with metastatic disease, treatment options are limited to cytotoxic chemotherapy or enrollment in clinical trials. To date, no clinical trial has demonstrated efficacy in the advanced or metastatic setting^5–7^, underscoring the urgent need for new therapeutic strategies.

A major genetic driver of MPNST pathogenesis is loss or inactivation of the *NF1* gene, which encodes neurofibromin, a negative regulator of RAS signaling. While *NF1* loss promotes benign tumor formation, it alone is insufficient for malignant transformation^8^. In genetically engineered mouse models *NF1* deletion in Schwann cell precursors leads to benign plexiform neurofibromas^9–11^, whereas additional genetic alterations, most commonly in *TP53*, *CDKN2A*, *EED*, or *SUZ12*, are required for malignant progression^12–15^. Among these, *SUZ12* and *EED* encode essential components of the Polycomb repressive complex 2 (PRC2), which catalyzes histone H3 lysine 27 trimethylation (H3K27me3) to silence developmental and lineage-specific genes^16^. Loss of PRC2 function occurs in most MPNSTs, with *SUZ12* and *EED* mutations present in approximately 56% and 32% of cases, respectively^17^. PRC2-deficient MPNSTs exhibit more aggressive clinical behavior and worse prognosis compared to PRC2-intact MPNSTs, particularly in NF1-associated cases^18–21^. Despite this, the downstream chromatin, transcriptional, and signaling consequences of PRC2 loss remain incompletely understood. Because loss-of-function mutations such as PRC2 deficiency are difficult to target directly, uncovering secondary, therapeutically actionable pathways downstream of epigenetic disruption is critical.

Insulin-like growth factor 2 (IGF2) is a fetal growth factor that plays a central role in prenatal growth and development^22^. In mice, loss of paternal Igf2 results in intrauterine growth restriction^23^, while IGF2 overexpression leads to increased body size^24^. IGF2 acts primarily through the IGF1 receptor (IGF1R) and an alternatively spliced isoform of the insulin receptor (IR-A). Additionally, its levels are tightly controlled by the mannose-6-phosphate/IGF2 receptor (M6P/IGF2R), which mediates IGF2 degradation^22,25^. Dysregulation of this pathway, through loss of imprinting, elevated IGF2 expression or altered receptor balance, has been implicated in multiple cancers where it promotes cell proliferation and survival^26,27^. Given PRC2’s role in developmental gene silencing, we reasoned that its loss might lead to inappropriate reactivation of fetal growth programs, including IGF2 signaling, to drive malignant growth.

To test this hypothesis, we performed integrative epigenomic, transcriptomic, and proteomic profiling of MPNST cell lines and primary tumors, comparing PRC2-intact and PRC2-deficient contexts. This genome-wide analysis revealed IGF2 among the top PRC2-regulated genes in MPNST. We showed that IGF2 is a direct and reversible target of H3K27me3-dependent repression and that PRC2-deficient MPNST cells acquire selective dependency on IGF2 signaling for growth. Together, these findings reveal a mechanism by which loss of PRC2 reactivates a fetal growth factor program to sustain malignant proliferation and identify IGF2 signaling as a potential therapeutic vulnerability in this aggressive tumor subtype.

## Results

### Epigenomic profiling reveals coordinated loss of H3K27me3 and gain of active chromatin marks in PRC2-deficient MPNST cells

To elucidate the effect of SUZ12 or EED loss on the chromatin landscape in MPNST, we analyzed a panel of MPNST cell lines (**Figure 1A**) by global histone post-translational modification (PTM) mass spectrometry (MS). In SUZ12-and EED-mutant cell lines, H3K27 methylations (H3K27me1, me2, me3, and their combinations with H3K36 methylations) were markedly reduced (**Figure 1B-C**), consistent with a PRC2-deficient state. In agreement with our previous work using FFPE tumors and a smaller panel of cell lines^28^, PRC2-deficient cells also displayed increased levels of activating histone PTMs, including H3K36 methylations (H3K36me1, H3K36me2, H3K36me3) and diverse combinations of H4 acetylations (e.g., H4K4acK8acK12acK16ac) (**Figure 1B**). H3K27ac showed a modest, but non-significant increase in PRC2-deficient cells (**Figure 1C**).

**Figure 1.**
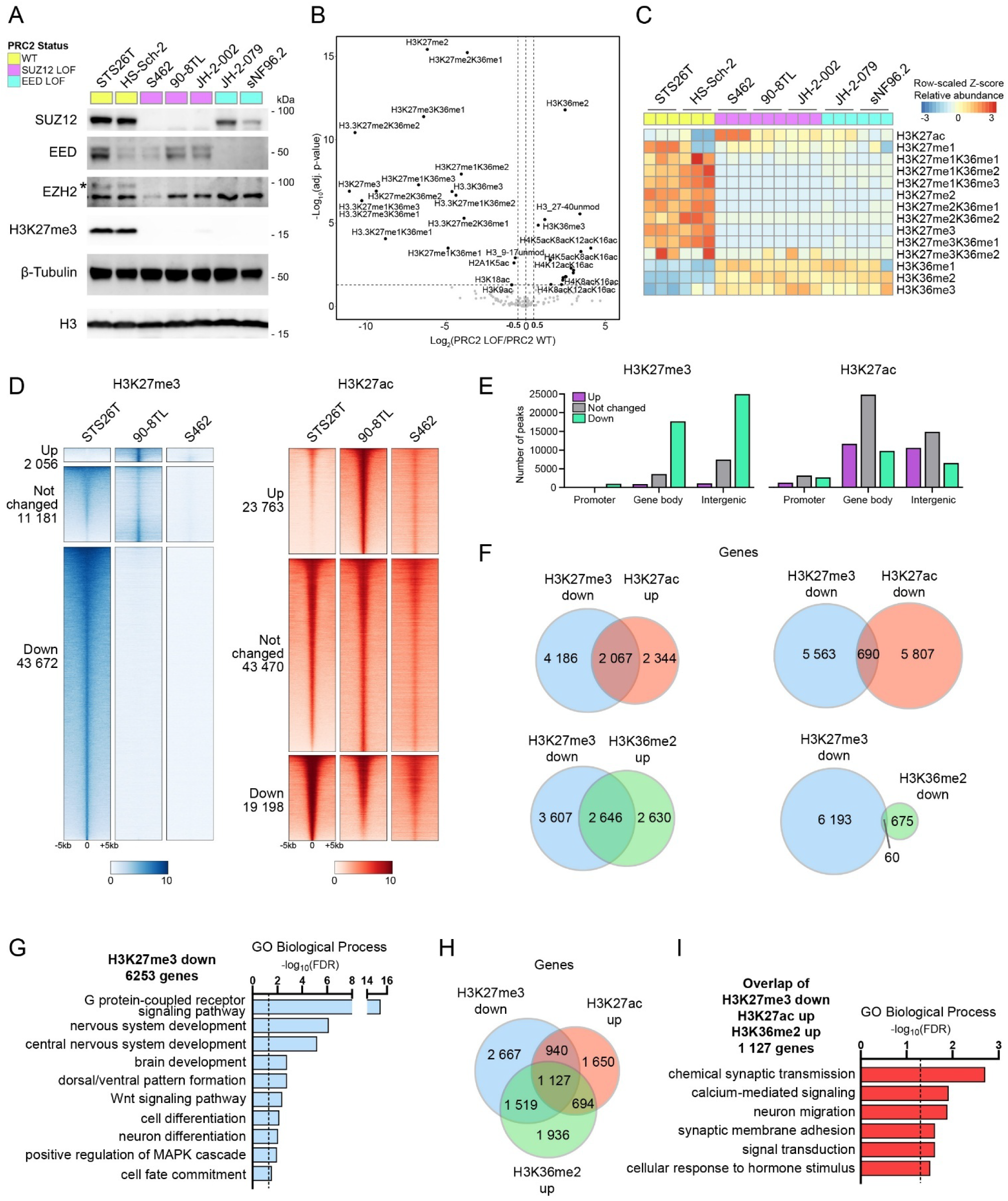
Epigenomic profiling reveals loss of H3K27me3 and gain of active chromatin marks in PRC2-deficient MPNST cells. **(A)** Levels of SUZ12, EED, EZH2 and H3K27me3 in PRC2-intact (WT) and PRC2-deficient (LOF) MPNST cell lines. **(B)** Significantly altered histone PTMs in PRC2-deficient cells detected by global histone PTM MS analysis (adj. p-value <0.05 and log2FC <-0.5 or >0.5). **(C)** Relative abundance of histone PTMs within the H3 27–40 peptide region from the MS data. Data represent three replicates per cell line. **(D)** H3K27me3 and H3K27ac ChIP-seq peaks (±5 kb) clustered as upregulated (adj. P < 0.05, log₂FC > 1), downregulated (adj. P < 0.05, log₂FC < -1), or unchanged in PRC2-deficient 90-8TL cells relative to PRC2-intact STS26T cells. The same peak regions are shown for PRC2-deficient S462 cells. **(E)** Genomic distribution of H3K27me3 and H3K27ac peak clusters from (D). **(F)** Overlap of genes with downregulated H3K27me3 and genes with up- or downregulated H3K27ac or H3K36me2. **(G)** Significantly enriched Gene Ontology (GO) Biological Processes (FDR < 0.05) for genes with downregulated H3K27me3. **(H)** Overlap of genes with downregulated H3K27me3 and upregulated H3K27ac and H3K36me2. **(I)** Significantly enriched GO Biological Processes (FDR < 0.05) for the overlapping genes in (H).

To map these histone PTMs genome-wide, we performed ChIP-seq for H3K27me3, H3K27ac, and H3K36me2 in PRC2-intact (STS26T) and PRC2-deficient (90-8TL and S462) cell lines. Consistent with the global histone PTM MS results, a large fraction of H3K27me3 peaks in STS26T cells (43,672 peaks) were significantly reduced in PRC2-deficient cells (adj. p < 0.05, log₂FC < -1) (**Figure 1D, Supplementary Fig. S1A**). In contrast, H3K27ac showed a similar number of up- and downregulated peaks (23,763 and 19,198 peaks, respectively) (adj. p < 0.05, |log₂FC| > 1) (**Figure 1D, Supplementary Fig. S1A**). Most downregulated H3K27me3 peaks were located at gene bodies (exons and introns) and intergenic regions, whereas a larger fraction of altered H3K27ac peaks were more frequently found at promoters (**Figure 1E**). Given the broad genomic distribution of H3K36me2 across both intergenic regions and gene bodies, with enrichment that scales with transcriptional activity^29,30^, we restricted our H3K36me2 analysis to gene-body read counts. In line with the MS data, most H3K36me2 changes in PRC2-deficient cells were increases (5,276 genes) rather than decreases (735 genes) (adj. p < 0.05, |log₂FC| > 1) (**Supplementary Fig. S1B**).

Next, we annotated the H3K27me3 and H3K27ac peaks to the nearest transcription start site (TSS) to identify potentially regulated genes. To avoid overestimating peak-gene associations from distal regions, we limited assignments to those within 10 kb of the TSS. Approximately one-third of genes with downregulated H3K27me3 (2,067 of 6,253) also showed upregulated H3K27ac, whereas overlap with downregulated H3K27ac was smaller (690 genes) (**Figure 1F**). A similar trend was observed for H3K36me2, where genes with reduced H3K27me3 predominantly showed increased H3K36me2 signal (2,646 genes upregulated vs. 60 downregulated) (**Figure 1F**).

To gain insight into the biological processes associated with these histone PTM changes, we performed Gene Ontology (GO) enrichment analysis of differentially modified gene groups, focusing on biological processes. Genes with downregulated H3K27me3 were enriched for differentiation and developmental pathways, including cell and neuron differentiation, nervous system and brain development, dorsal-ventral pattern formation, and cell fate commitment (**Figure 1G**). Interestingly, this group also included genes involved in the positive regulation of the MAPK cascade (**Figure 1G**).

GO analysis of genes with upregulated H3K27ac revealed similar enrichment for developmental and differentiation pathways (**Supplementary Fig. S1C**), whereas genes with downregulated H3K27ac were enriched for processes related to DNA repair, chromatin remodeling, splicing, and translation (**Supplementary Fig. S1C**). Genes with upregulated H3K36me2 were associated with pathways involved in synaptic transmission and membrane potential, as well as immune and inflammatory responses, including cell chemotaxis and positive regulation of immune response (**Supplementary Fig. S1D**). Notably, 1,127 genes exhibited coordinated changes across all three histone PTMs; loss of H3K27me3 together with gain of H3K27ac and H3K36me2 (**Figure 1H**) and were enriched for processes such as calcium-mediated signaling, neuron migration, and synaptic membrane adhesion (**Figure 1I**).

Together, these results suggest that loss of the repressive H3K27me3 mark near TSSs can promote acquisition of activating histone marks, such as H3K27ac and H3K36me2, at genes involved in differentiation, development, and nervous system -specific programs. While some genes show coordinated remodeling of all three marks, others display selective modification, suggesting that these histone PTMs contribute to both shared and distinct regulatory functions in PRC2-deficient MPNST cells.

### PRC2 loss promotes IGF2 transcriptional activation via loss of H3K27me3 and gain of H3K27ac

To investigate the transcriptional consequences of PRC2 loss, we performed RNA-seq across the panel of MPNST cell lines. Differential expression analysis revealed that 3,593 genes were significantly upregulated and 1,953 genes downregulated in PRC2-deficient cells (S462, 90-8TL, JH-2-002, JH-2-079, sNF96.2) compared to PRC2-intact cells (STS26T and HS-Sch-2) (adj. p-value < 0.05, |log₂FC| > 1) (**Figure 2A, Supplementary Fig. S2A–B**). Gene Ontology (GO) analysis of upregulated genes showed enrichment for processes such as cell differentiation, nervous system development, and positive regulation of MAPK signaling (**Figure 2B**), consistent with pathways associated with loss of H3K27me3 (**Figure 1G**). In contrast, downregulated genes were enriched for biological processes related to antigen processing and presentation via MHC class II, extracellular matrix disassembly, and inflammatory and immune responses (**Figure 2B**).

**Figure 2.**
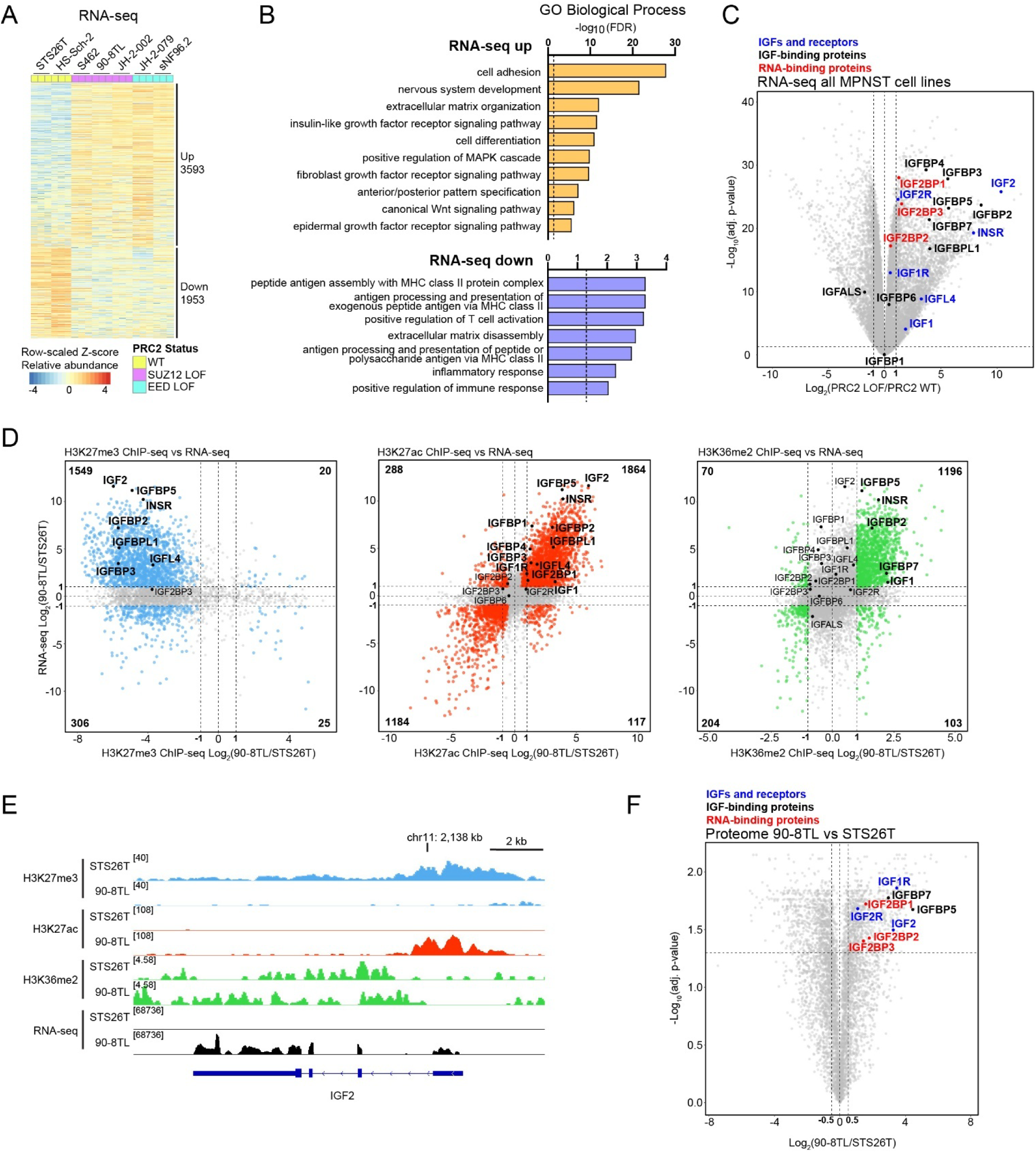
Multi-omics integration identifies activation of IGF2 signaling in PRC2-deficient MPNST cells. **(A)** Differentially expressed genes in PRC2-deficient (LOF) versus PRC2-intact (WT) MPNST cell lines identified by RNA-seq in three biological replicates (adj. p-value < 0.05 and |log_2_FC| > 1). **(B)** Significantly enriched Gene Ontology (GO) Biological Processes (FDR < 0.05) for the up- and downregulated genes from (A). **(C)** Volcano plot highlighting IGF2-signaling -related genes from the RNA-seq comparison between PRC2-deficient and PRC2-intact cells. **(D)** Scatter plots showing the relationship between transcriptional and chromatin changes. The Y-axis represents RNA-seq log₂FC (90-8TL vs STS26T), and the X-axis represents ChIP-seq log₂FC for H3K27me3 (left), H3K27ac (middle), and H3K36me2 (right). Colored points indicate genes significantly altered in both datasets (adj. p-value <0.05 and log2FC <-1 or >1). Numbers in corners denote the number of genes in that category. IGF2-signaling -related genes are highlighted. **(E)** Genome browser (IGV) tracks showing H3K27me3, H3K27ac, H3K36me2, and RNA-seq signal at the *IGF2* locus in PRC2-intact (STS26T) and PRC2-deficient (90-8TL) cells. **(F)** Volcano plot of quantitative proteomics comparing PRC2-deficient 90-8TL and PRC2-intact STS26T cells, highlighting IGF2-signaling -related proteins.

Among the upregulated pathways, the insulin-like growth factor (IGF) signaling cascade emerged as one of the most significantly enriched (**Figure 2B**). Insulin-like growth factor 2 (IGF2) was one of the most highly upregulated genes in PRC2-deficient cells (**Figure 2C**), whereas related ligands IGF1 and IGF-like family member 4 (IGFL4) showed more modest increases. Several genes encoding IGF2-interacting or regulatory proteins were also upregulated, including IGF2 receptors (IGF2R and INSR), IGF2 mRNA-binding proteins (IGF2BP1 and IGF2BP3), and multiple IGF-binding proteins (IGFBP2, IGFBP3, IGFBP4, IGFBP5, IGFBP7, IGFBPL1). These findings suggest a coordinated activation of the IGF2 signaling network following PRC2 loss.

To determine how PRC2 loss -associated histone changes contribute to this transcriptional activation, we integrated the RNA-seq and ChIP-seq datasets. Most genes that lost H3K27me3 showed increased mRNA expression (1,549 genes), whereas H3K27ac and H3K36me2 were positively correlated with gene expression: genes with increased H3K27ac (1,864 genes) or H3K36me2 (1,196 genes) were typically upregulated, while those with decreased signal were downregulated (1,184 and 204 genes, respectively) (**Figure 2D, Supplementary Fig. S2D**). IGF2 displayed one of the strongest decreases in H3K27me3 together with a marked increase in H3K27ac and a modest, nonsignificant gain in H3K36me2 (**Figure 2D-E**). Quantitative proteomics further confirmed elevated IGF2 protein levels together with increased abundance of multiple IGF2 signaling components (**Figure 2F, Supplementary Fig. S2E-F**).

Together, these results indicate that PRC2 loss drives coordinated transcriptional and proteomic activation of IGF2 signaling in MPNST cells. IGF2 derepression is primarily associated with loss of the repressive H3K27me3 mark and concomitant gain of H3K27ac, while H3K36me2 appears to play a lesser role in regulating IGF2 expression.

### Restoration of SUZ12 in PRC2-deficient MPNST cells re-establishes H3K27me3 and reverses IGF2 expression

To validate the histone PTM, transcriptomic, and proteomic changes in an isogenic setting, we generated a doxycycline (Dox)-inducible SUZ12 re-expression model using the PRC2-deficient 90-8TL cells. After 7 days of Dox induction, H3K27me3 levels were restored to levels comparable to PRC2-intact cell lines (**Figure 3A, Supplementary Fig. S3A**). ChIP-seq for H3K27me3 and H3K27ac confirmed global remodeling of chromatin states following SUZ12 restoration. As expected, H3K27me3 peaks were predominantly gained, with 25,943 sites significantly upregulated upon Dox treatment (adj. p < 0.05, log₂FC > 1) (**Figure 3B-C**). In contrast, H3K27ac peaks showed a nearly equal number of gains and losses (6,200 and 6,196 peaks, respectively) (adj. p < 0.05, |log₂FC| > 1) (**Figure 3B-C, Supplementary Fig. S3B**). Newly gained H3K27me3 sites were mainly located at intergenic and gene body regions (exons and introns), whereas altered H3K27ac peaks were more frequently found at promoters and gene bodies (**Figure 3D**).

**Figure 3.**
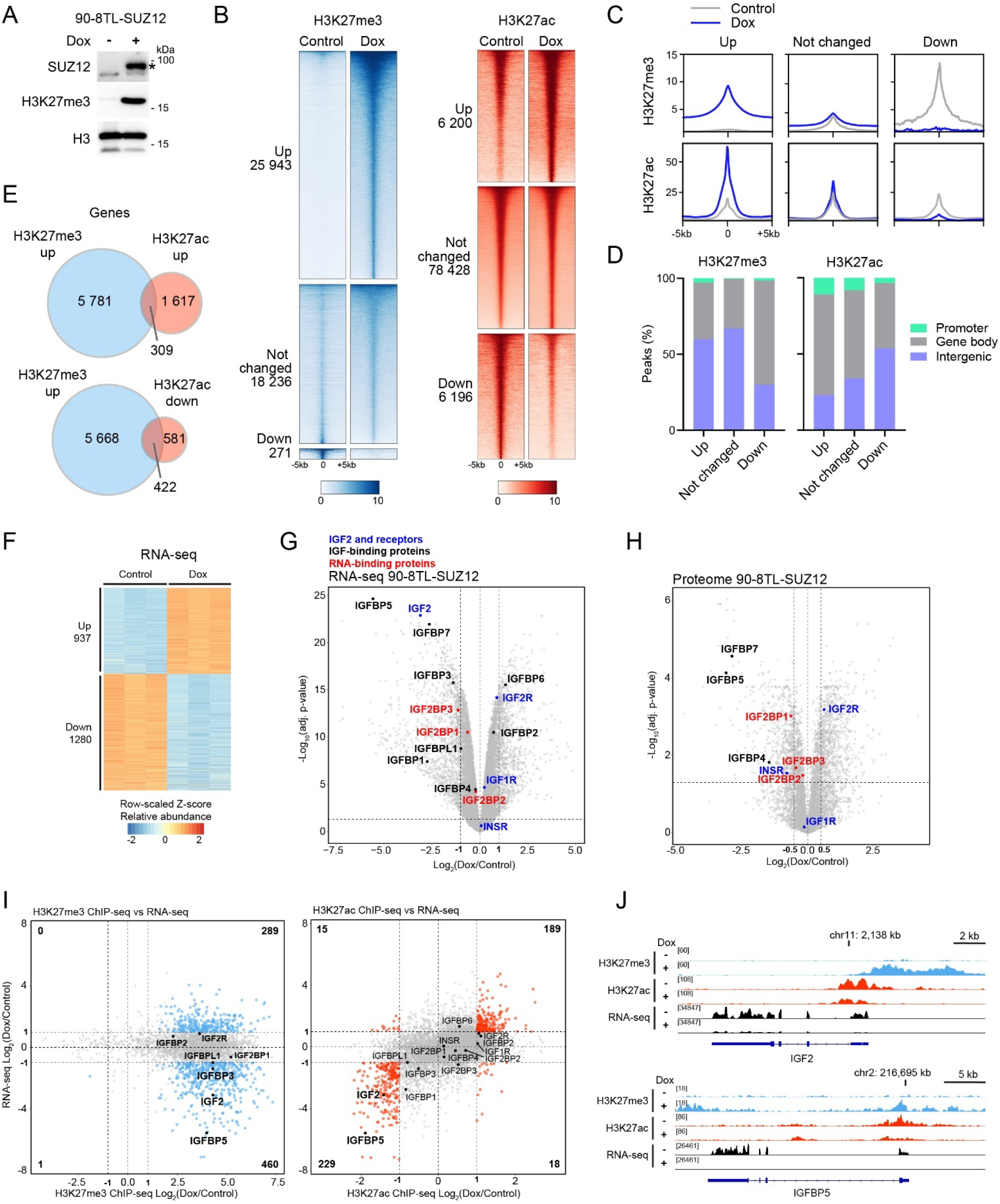
Restoration of SUZ12 in PRC2-deficient MPNST cells re-establishes H3K27me3 and reverses IGF2 expression. **(A)** Levels of SUZ12 and H3K27me3 in doxycycline (Dox) -inducible 90-8TL-SUZ12 cells with or without Dox treatment for 7 days. Asterix depicts specific SUZ12 band. **(B)** H3K27me3 and H3K27ac ChIP-seq peaks (±5 kb) clustered as upregulated (adj. p-value < 0.05, log₂FC > 1), downregulated (adj. p-value < 0.05, log₂FC < -1), or unchanged in SUZ12-restored cells (Dox) relative to PRC2-deficient control cells. **(C)** Line plots showing average ChIP-seq peak profiles for each cluster from (B). **(D)** Genomic distribution of H3K27me3 and H3K27ac peak clusters from (B). **(E)** Overlap of genes with upregulated H3K27me3 and genes with up- or downregulated H3K27ac. **(F)** Differentially expressed genes in Dox-treated versus control cells identified by RNA-seq (adj. p-value <0.05 and log2FC < -1 or > 1). **(G)** Volcano plot highlighting IGF2-signaling -related genes from the RNA-seq comparison between Dox and control treatment. **(H)** Volcano plot highlighting IGF2-signaling-related proteins from quantitative proteomics of Dox versus control treatment. **(I)** Scatter plots showing the relationship between transcriptional and chromatin changes. The Y-axis represents RNA-seq log₂FC (Dox vs Control), and the X-axis represents ChIP-seq log₂FC for H3K27me3 (left) and H3K27ac (right). Colored points indicate genes significantly altered in both datasets (adj. p-value <0.05 and log2FC < -1 or > 1). Numbers in corners denote the number of genes in that category. IGF2-signaling -related genes are highlighted. **(J)** Genome browser (IGV) tracks showing H3K27me3, H3K27ac, and RNA-seq signal at the *IGF2* and *IGFBP5* loci in dox and control treated cells.

Annotation of peaks within 10 kb of TSSs revealed that a subset of genes (422 of 6,090) with increased H3K27me3 also showed reduced H3K27ac, while a smaller group (309 genes) gained both marks (**Figure 3E**). GO analysis of genes with gained H3K27me3 indicated enrichment for pathways related to cell development, differentiation, cell fate commitment, and MAPK signaling, consistent with data from the unmodified MPNST cell lines. Genes with reduced H3K27ac similarly included pathways involved in differentiation-related processes (**Supplementary Fig. S3C**).

To examine the transcriptional consequences of PRC2 restoration, we performed RNA-seq at the 7-day Dox induction time point. SUZ12 re-expression resulted in 937 upregulated and 1,280 downregulated genes (adj. p < 0.05, |log₂FC| > 1) (**Figure 3F, Supplementary Fig. S3D**). Consistent with the ChIP-seq findings, downregulated genes were enriched for differentiation and developmental processes, whereas upregulated genes were associated with cell adhesion and inflammatory pathways (**Supplementary Fig. S3E**). Importantly, IGF2 was among the most strongly downregulated genes, accompanied by decreased expression of several IGF2-related components, including IGFBP5, IGFBP7, and IGF2BP3 (**Figure 3G**). Proteomic analysis showed a similar trend, although IGF2 and several IGFBPs were not detected, possibly reflecting their low intracellular abundance due to being secreted into the medium. (**Figure 3H, Supplementary Fig. S3F**).

Integration of RNA-seq and ChIP-seq datasets revealed that genes gaining H3K27me3 were more often repressed (460 genes) than activated (289 genes) following SUZ12 restoration (**Figure 3I, Supplementary Fig. S3G**). In contrast, changes in H3K27ac showed a clear positive correlation with gene expression, where genes with increased H3K27ac (189 genes) were upregulated and those with decreased H3K27ac (229 genes) were downregulated. Notably, IGF2 and IGFBP5 showed two of the strongest examples of concurrent H3K27me3 gain and H3K27ac loss after Dox treatment (**Figure 3I– J**).

Together, these results demonstrate that SUZ12 restoration reverses IGF2 expression by re-establishing H3K27me3 and reducing H3K27ac levels in PRC2-deficient MPNST cells. Thus, PRC2 loss is both necessary and sufficient to drive IGF2 activation, directly linking PRC2 deficiency to aberrant growth signaling in MPNST.

### Restoration of SUZ12 in PRC2-deficient MPNST cells reduces IGF2 secretion, and IGF2 knockdown selectively impairs proliferation in the PRC2-deficient state

Because IGF2 acts as a secreted growth factor, we next asked whether SUZ12 restoration affects its secretion and functional impact on proliferation. We therefore performed IGF2 enzyme-linked immunosorbent assay (ELISA) from medium conditioned with Doxycycline-inducible SUZ12-restoration cells. After 72 hours of culture, PRC2-deficient cells secreted an average of 2.4 ng/ml IGF2 per million cells (**Figure 4A**). Dox-induced SUZ12 restoration decreased IGF2 secretion by more than 50%. Knockdown of IGF2 in either condition reduced IGF2 levels to near background.

**Figure 4.**
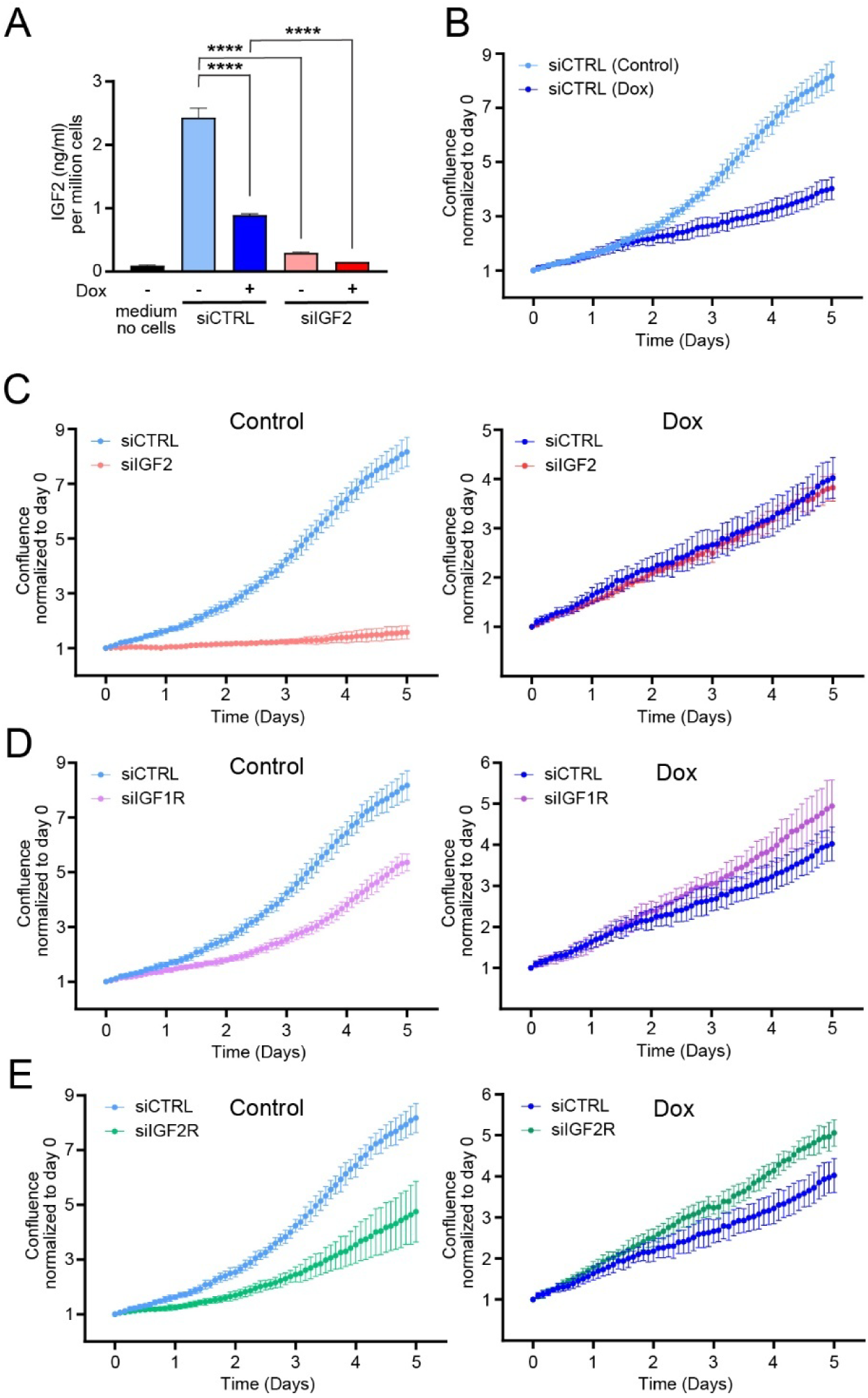
Restoration of SUZ12 in PRC2-deficient MPNST cells reduces IGF2 secretion, and IGF2 knockdown selectively impairs proliferation in the PRC2-deficient state. **(A)** IGF2 ELISA from conditioned medium of PRC2-deficient 90-8TL (Control) or SUZ12-restored (Dox-treated) cells transfected with non-targeting control siRNA (siCTRL) or IGF2 siRNA (siIGF2). Two-way ANOVA; **** adj. p-value < 0.0001. **(B)** Normalized cell confluence over five days for control and Dox-treated cells transfected with siCTRL. Normalized cell confluence over five days for control and Dox-treated cells transfected with siRNAs targeting **(C)** IGF2, **(D)** IGF1R, or **(E)** IGF2R. Data represent mean ± SD from three biological replicates.

Next, we used live-cell imaging to examine the impact of IGF2 on proliferation. SUZ12 restoration alone markedly reduced cell growth (**Figure 4B**). Interestingly, IGF2 knockdown impaired proliferation only in PRC2-deficient cells, whereas SUZ12-restored cells were unaffected (**Figure 4C**). To test whether this dependency extends to IGF2-binding receptors, we performed siRNA-mediated knockdowns of IGF1R and IGF2R. In PRC2-deficient cells, knockdown of either receptor reduced proliferation, whereas in SUZ12-restored cells, knockdown slightly increased proliferation (**Figure 4D-E**).

These results demonstrate that secreted IGF2 levels are directly regulated by SUZ12 and that PRC2-deficient MPNST cells are specifically dependent on IGF2 for growth. Both IGF1R and IGF2R mediated this effect, underscoring the functional importance of the IGF2-mediated signaling in PRC2-deficient state.

### PRC2 loss in MPNSTs is associated with altered histone modification patterns and upregulation of the IGF2-IGF2BP axis

To assess the clinical relevance of IGF2 upregulation, we analyzed the total proteome of a panel of human MPNSTs and plexiform neurofibromas (PNs). Tumor PRC2 status was verified by mass spectrometry-based histone PTM analysis. Four tumors were classified as PRC2-deficient, showing patterns similar to PRC2-deficient cell lines: downregulation of H3K27 methylations, upregulation of H3K36 methylations, increased combinatorial H4 acetylations (e.g., H4K5acK8acK16ac), and a consistent but non-significant trend toward increased H3K27ac (**Figure 5A-B, Supplementary Fig. S4A**).

**Figure 5.**
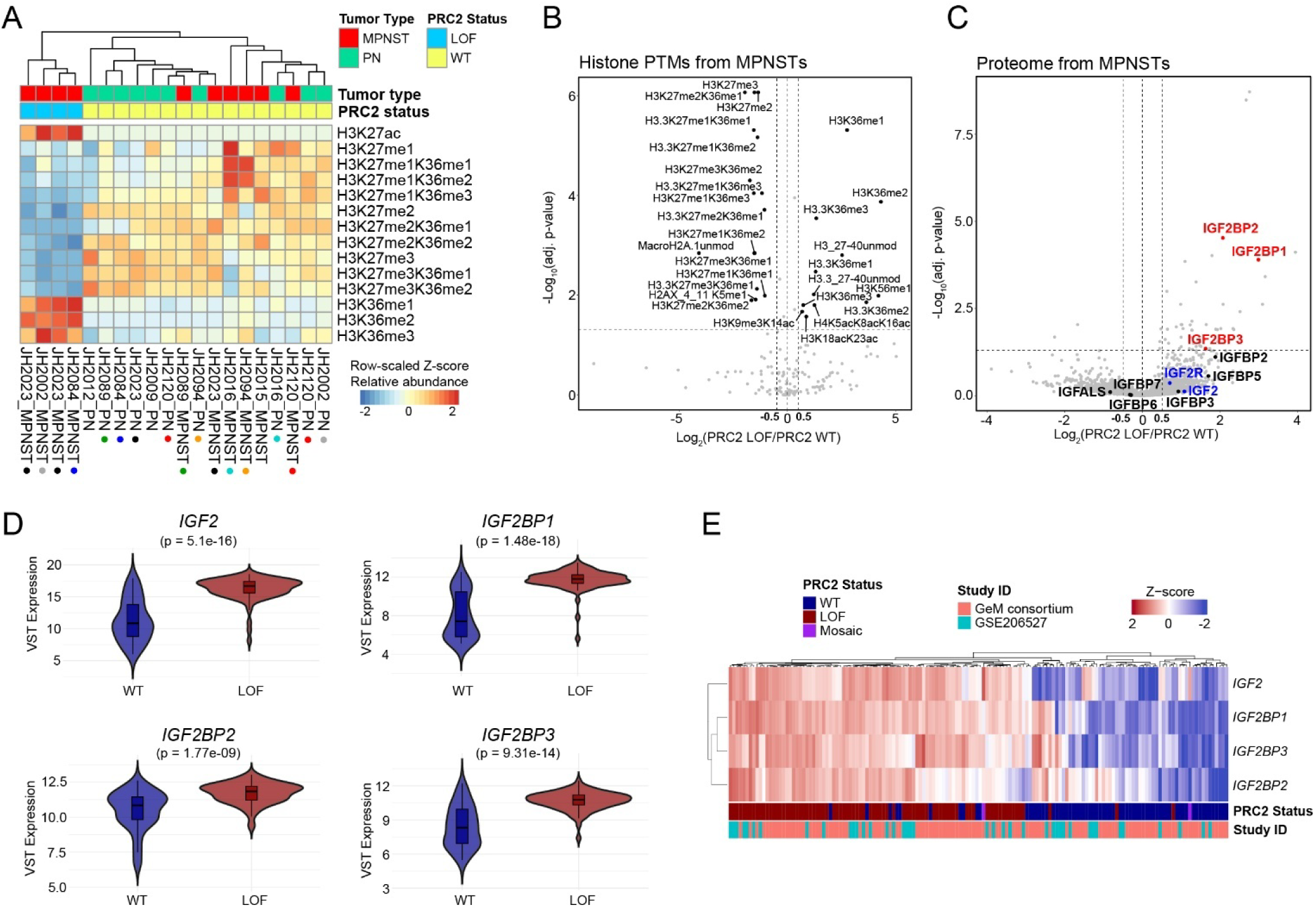
PRC2 loss in MPNSTs is associated with altered histone modification patterns and upregulation of the IGF2-IGF2BP axis. **(A)** Relative abundance of histone PTMs within the H3 27-40 peptide region from human PRC2-intact (WT) or PRC2-deficient (LOF) MPNSTs and PNs detected by global histone PTM MS analysis. Samples are ordered by hierarchical clustering. Colored dots below the heatmap indicate paired samples from the same patient. **(B)** Significantly altered histone PTMs in PRC2-deficient versus PRC2-intact MPNSTs (adj. p < 0.05 and log₂FC < -0.5 or > 0.5). **(C)** Volcano plot highlighting IGF2-signaling -related proteins from quantitative proteomics comparing PRC2-deficient and PRC2-intact MPNSTs. **(D)** Violin plots showing average mRNA expression of *IGF2*, *IGF2BP1*, *IGF2BP2*, and *IGF2BP3* in PRC2-WT and -LOF tumors from the MPNST cohorts (GeM and GSE206527). Statistical significance was assessed by Wilcoxon test. **(E)** Hierarchically clustered heatmap showing relative expression of *IGF2*, *IGF2BP1*, *IGF2BP2*, and *IGF2BP3* in individual samples from the MPNST cohorts.

Comparative proteome analysis of PRC2-deficient (n=4) versus PRC2-intact (n=7) MPNSTs revealed significant upregulation of IGF2BP1-3 proteins in PRC2-deficient tumors (adjusted p < 0.05, |log₂FC| > 0.5) (**Figure 5C**). IGF2 and several IGFBPs showed modest upregulation but did not reach significance, likely due to their secretory nature and the limited sample size. Notably, the tumor panel included two PRC2-deficient MPNSTs, one PRC2-intact MPNST, and one PN from the same patient. Within this patient, IGF2 and IGF2BP expression was elevated specifically in the PRC2-deficient tumors (**Supplementary Fig. S4B**).

Because IGF2 was not significantly upregulated in the proteomics dataset, we next analyzed publicly available transcriptomic data from 150 human MPNSTs (Genomics of MPNST [GeM] consortium and GSE206527 datasets)^18,31^. In these datasets, IGF2 and IGF2BP1-3 were all significantly upregulated in PRC2-deficient tumors (p = 5.1e-16, 1.48e-18, 1.77e-9, and 9.31e-14, respectively; **Figure 5D**). Hierarchical clustering based on IGF2 and IGF2BP1-3 expression largely segregated tumors by PRC2 status (**Figure 5E**).

In contrast, other IGF ligands, IGF1 and IGFL4, were moderately but significantly downregulated in PRC2-deficient tumors (p = 0.0161 and 0.001, respectively; **Supplementary Fig. S4C**). Among IGF-binding receptors, IGF1R was significantly upregulated (p = 3.95e-5), whereas Insulin Receptor (INSR) and IGF2R showed no significant difference (**Supplementary Fig. S4D**). Analysis of IGFBPs revealed significant upregulation of IGFBP2 and IGFBPL1 (p = 6.28e-9 and 1.87e-10, respectively), while IGFBP4, IGFBP6, and IGFBP7 were downregulated (p = 0.0166, 9.47e-9, and 1.08e-6, respectively; **Supplementary Fig. S4E**). IGFBP1, IGFBP3, and IGFBP5 were unchanged.

Taken together, these results demonstrate that both IGF2 and IGF2BP1-3 are significantly upregulated in clinical PRC2-deficient MPNSTs. The data link PRC2 loss to the activation of the IGF2-IGF2BP axis and suggests that this pathway may contribute to the more aggressive phenotype of PRC2-deficient tumors, highlighting IGF2 and its regulatory proteins as potential biomarkers and therapeutic targets.

## Discussion

In this study, we demonstrate that PRC2 loss in MPNST induces coordinated chromatin remodeling characterized by loss of the repressive mark H3K27me3 and acquisition of activating marks, particularly H3K27ac and H3K36me2. Integration of epigenomic, transcriptomic, and proteomic datasets revealed that this chromatin reprogramming activates a fetal-like growth signature centered on IGF2 and its post-transcriptional regulators IGF2BP1-3, linking PRC2 deficiency to aberrant growth factor signaling. Functional assays further establish that PRC2-deficient MPNST cells are selectively dependent on IGF2 for proliferation, highlighting the IGF2-IGF2BP axis as a key effector and potential vulnerability of this epigenetic subtype.

Loss of H3K27me3 is the hallmark of PRC2 inactivation, yet its downstream chromatin consequences in MPNST have not been comprehensively defined. Our histone PTM mass spectrometry and ChIP-seq analyses reveal a broad shift from repressive to active chromatin states, consistent with prior reports of PRC2-deficient MPNST by us and others^28,32^. The preferential gain of H3K36 methylation, H3K27 acetylation and combinatorial H4 acetylations supports a global relaxation of chromatin compaction and enhanced transcriptional competence. Importantly, we show that a subset of loci, particularly those related to neuronal signaling and development, such as chemical synaptic tansmission, calcium-mediated signaling, and neuron migration, undergo coordinated loss of H3K27me3 and gain of both H3K27ac and H3K36me2. This underscores a highly ordered reprogramming rather than stochastic derepression.

These chromatin changes correspond closely with transcriptional activation of developmental and growth pathways. The enrichment of nervous system and cell fate specification genes among H3K27me3-lost and H3K27ac-gained loci suggests that PRC2 loss reawakens neural crest-like programs. This is consistent with the developmental origin of Schwann cells and may contribute to the plasticity and aggressive behavior of PRC2-deficient MPNSTs.

Among the genes most strongly derepressed by PRC2 loss was IGF2, a well-established fetal growth factor normally silenced postnatally by epigenetic imprinting^22^. It is frequently upregulated in cancer and promotes cell proliferation and survival^26,27^. A previous study identified IGF2 as one of the overexpressed genes in PRC2-deficient MPNST cells and showed that its promoter gained H3K27ac and H3K4me3 while losing H3K27me3, SUZ12 and EZH2, reversible by SUZ12 re-expression^17^.

Our integrative, unbiased analyses support and extend these findings, identifying IGF2 among the top PRC2-regulated genes with strong activation accompanied by pronounced loss of H3K27me3 and gains of H3K27ac at its promoter, consistent with direct transcriptional derepression. The modest increase H3K36me2 increase at the IGF2 locus suggests that IGF2 is not directly under the regulation of H3K36me2. However, several IGF-related genes, IGF1, INSR, IGFBP2, IGFBP5, and IGFBP7, showed increased gene-body H3K36me2 in PRC2-deficient cells, implying a supporting role in the wider signaling network. Given that H3K36me3 is more strongly associated with active transcription^29,30^, future studies profiling this mark may reveal more pronounced effects. Importantly, restoration of SUZ12 re-established H3K27me3 at the IGF2 locus, reduced H3K27ac, and sharply downregulated IGF2 mRNA and protein, confirming that PRC2 activity is both necessary and sufficient for IGF2 silencing.

These results reveal that loss of PRC2 can override imprinting-like repression at the IGF2 locus in a somatic tumor context. The ability to reverse IGF2 activation through SUZ12 restoration provides a mechanistic link between PRC2 deficiency and aberrant growth factor signaling in MPNST.

The functional role of IGF2 in PRC2-deficient MPNSTs had not been defined previously. Our functional assays establish the biological relevance of IGF2 activation. PRC2-deficient cells secreted high levels of IGF2, and SUZ12 re-expression reduced both IGF2 secretion and proliferation.

Moreover, IGF2 knockdown impaired proliferation only in the PRC2-deficient state, demonstrating a context-specific dependency. Similarly, both IGF1R and IGF2R knockdown individually impaired proliferation only in PRC2-deficient cells, though not to the same extent as IGF2 knockdown, suggesting that IGF2 signals through multiple receptors to sustain growth.

In mice, lowering *Igf2r* expression results in elevated circulating IGF2 and increased growth, consistent with its role in antagonizing IGF2 signaling^22^. However, *IGF2R* knockdown in PRC2-deficient human MPNST cells reduced proliferation, suggesting that it plays a positive role in IGF2 signaling under these conditions. This effect was reversed in SUZ12-restored cells for both IGF1R and IGF2R, where knockdown of either receptor slightly increased growth, indicating that they antagonize IGF2 signaling after PRC2 restoration. Taken together, these results suggest that PRC2 loss establishes an autocrine IGF2 signaling loop that drives proliferation, a mechanism reminiscent of developmental growth programs reactivated in cancer.

Proteomic profiling of human MPNSTs confirmed that PRC2-deficient tumors exhibit histone PTM patterns consistent with cell line models. Notably, IGF2BP1-3 were significantly upregulated in PRC2-deficient tumors, and transcriptomic data from large patient cohorts corroborated co-upregulation of *IGF2* and *IGF2BP1-3*. Given the established roles of IGF2BPs in stabilizing and enhancing translation of IGF2 mRNAs^33–35^, their induction likely amplifies IGF2 signaling and post-transcriptional control of growth-promoting genes. Thus, PRC2 loss appears to activate a coordinated IGF2-IGF2BP axis that may reinforce oncogenic programs and promote tumor aggressiveness. This axis could also serve as a molecular biomarker for PRC2-loss MPNSTs, with potential diagnostic or prognostic value.

Our findings support a model in which PRC2 loss reshapes the chromatin landscape to derepress fetal growth regulators, particularly *IGF2*, thereby establishing an autocrine signaling circuit that promotes proliferation. The resulting activation of the IGF2-IGF1R/IGF2R network may synergize with RAS pathway hyperactivation driven by *NF1* loss, further enhancing oncogenic signaling. These data suggest that targeting IGF2 signaling, through IGF1R inhibitors or emerging IGF2BP antagonists, could provide a therapeutic strategy for PRC2-deficient MPNSTs, a subgroup currently lacking effective targeted options. Moreover, our results underscore the reversibility of PRC2-loss -driven transcriptional programs, raising the possibility that pharmacologic repression of H3K27 acetylation could attenuate IGF2-driven growth.

While our data establish a causal link between PRC2 loss and *IGF2* activation, further studies are needed to elucidate how enhancer-promoter interactions and three-dimensional genome architecture contribute to *IGF2* derepression. *In vivo* validation will be critical to determine whether IGF2 dependency extends to PRC2-deficient MPNST tumors and whether inhibition of IGF2 signaling or chromatin regulators can achieve therapeutic benefit. Combinatorial receptor knockdown experiments may help identify the receptors mediating IGF2 signaling, providing insight into which receptor targets may be most effective for drug development.

Interestingly, IGF2 prohormones are secreted by sarcoma cells^36^ and were recently shown to exert the strongest effects on osteoblast differentiation and proliferation^37^. It remains to be examined whether these higher-molecular weight forms of IGF2 contribute to MPNST growth. Finally, determining whether IGF2BPs directly regulate *IGF2* mRNA stability, and whether their stabilization of other target RNAs intersects with PRC2-loss -induced transcriptional changes, may uncover novel post-transcriptional vulnerabilities.

In summary, this work identifies a direct link between PRC2 loss and IGF2-driven autocrine signaling, establishing the IGF2-IGF2BP axis as a defining feature and potential vulnerability of PRC2-deficient MPNSTs. These results open new avenues for therapeutic intervention in this aggressive tumor type and underscore the broader principle that chromatin-state alterations can create targetable growth dependencies.

## Methods

### Cell lines and culture conditions

sNF96.2 cells were obtained from the American Type Culture Collection (ATCC), and HS-Sch-2 cells were purchased from the RIKEN BioResource Research Center (Japan). JH-2-002 and JH-2-079 cells were obtained from the Johns Hopkins University NF1 Biospecimen Repository (courtesy of Dr. Christine Pratilas). 90-8TL and S462 cell lines were generously provided by Dr. Karen Cichowski (Dana-Farber Cancer Institute, Harvard Medical School), and STS26T cells were provided by Dr. Jeffrey Field (Perelman School of Medicine, University of Pennsylvania). All cell lines were maintained in Dulbecco’s Modified Eagle Medium (DMEM; Corning, 10-013-CV) supplemented with 10% fetal bovine serum (FBS; R&D Systems, S12450), 1% penicillin-streptomycin (Gibco, Thermo Fisher Scientific, 15140122), and 1% GlutaMAX™ (Gibco, Thermo Fisher Scientific, 35050061). Cells were routinely tested for mycoplasma contamination and authenticated by short tandem repeat (STR) profiling performed by the Genome Engineering and Stem Cell Center (GESC), Washington University in St. Louis. STR profiles are available upon request.

### Generation of doxycycline-inducible SUZ12 re-expression line

For construction of the doxycycline-inducible SUZ12 expression vector, the human *SUZ12* open reading frame (ORF) from the pCMV-HA-SUZ12 plasmid (Addgene, #24232; gift from Kristian Helin)^38^ was cloned into the piggyBac Xlone-puro backbone by replacing the *eGFP* sequence in Xlone-puro-eGFP (Addgene, #140027; gift from Xiaoping Bao). The integrity of the resulting Xlone-puro-SUZ12 construct was verified by Sanger sequencing.

For stable cell line generation, 90-8TL cells were co-transfected with Xlone-puro-SUZ12 and the Super PiggyBac Transposase Expression Vector (System Biosciences, #PB210PA-1) using Lipofectamine 3000 (Thermo Fisher Scientific) according to the manufacturer’s protocol. Three days after transfection, cells were selected with 1 µg/mL puromycin (Thermo Fisher Scientific, #A1113803) for three weeks to establish the TRE3G-SUZ12-90-8TL stable line.

### Western blotting

Approximately 1.5 × 10^6^ cells were washed with PBS, trypsinized, and trypsin was quenched with complete medium. Cells were collected by centrifugation, and the pellets were washed twice with PBS. Cell pellets were lysed in 150 µL Laemmli buffer (Bio-Rad, #610737) containing 2.5% β-mercaptoethanol, protease inhibitor cocktail (Sigma-Aldrich, #P8340), and 10 mM sodium butyrate (Sigma-Aldrich, #303410). Lysates were heated at 95°C for 10 min and sonicated twice with a probe sonicator (Fisherbrand, Thermo Fisher Scientific) for 10 s at 20% amplitude. Aliquots of 10-20 µL were loaded on 10% or 16% Novex WedgeWell Tris-Glycine gels together with Precision Plus All Blue or Kaleidoscope protein ladders (Bio-Rad). Gels were run for approximately 40 min at 150 V on a Novex Mini Gel Tank (Thermo Fisher Scientific) using Tris-Glycine running buffer (Bio-Rad). Proteins were transferred to nitrocellulose membranes for 1 h at 100 V in a cooled Bio-Rad transfer system using Tris-Glycine transfer buffer supplemented with 20% methanol.

Membranes were blocked in Intercept TBS Blocking Buffer (LI-COR) for 1 h at room temperature, then incubated overnight at 4°C with primary antibodies diluted in the same blocking buffer. Primary antibodies were from Cell Signaling Technology: anti-SUZ12 (#3737S), anti-EED (#85322S), anti-EZH2 (#5246S), anti-H3K27me3 (#9733S), anti-β-Tubulin (#86298), and anti-H3 (#9715). After three washes in TBS containing 0.1% Tween-20 (TBS-T), membranes were incubated for 1 h at room temperature in blocking buffer containing HRP-conjugated secondary antibody (Cell Signaling Technology). Blots were washed three additional times with TBS-T and developed using Amersham™ ECL Prime Western Blotting Detection Reagent (Cytiva). Signals were visualized on an Odyssey imaging system (LI-COR), and images were processed using ImageJ^39^.

### siRNA transfection and cell proliferation assays

TRE3G-SUZ12-90-8TL cells were cultured in standard growth medium supplemented with 1 µg/mL puromycin and 100 ng/mL doxycycline (Sigma-Aldrich, #D9891) or water as a control for 4 days. The medium was then replenished, and cells were cultured for an additional 3 days. After a total of 7 days of doxycycline induction, cells were seeded in 6-well plates and cultured for 24 h prior to transfection.

Cells were transfected with 20 nM ON-TARGETplus SMARTpool siRNAs (Dharmacon) targeting IGF2 (L-004093-00-0005), IGF1R (L-003012-00-0005), IGF2R (L-010601-00-0005), or non-targeting control siRNA (D-001810-10-20) using standard growth medium supplemented with 1 µg/mL puromycin and 100 ng/mL doxycycline (or water as control). Three days after transfection, cells were transferred to 96-well plates at a density of 3,000 cells per well in three biological replicates. The medium was supplemented with 1 µg/mL puromycin, 100 ng/mL doxycycline (or water as control), and 25 nM YOYO-1 dye (Biotium).

Cells were allowed to attach for 24 h, after which cell confluence and YOYO-1 green fluorescence were monitored for 5 days using live-cell imaging on the IncuCyte S3 system (Sartorius). Cell confluence was quantified automatically using the IncuCyte S3 Image Analysis Software. Briefly, phase-contrast images were analyzed with a minimum area filter of 200 µm², segmentation adjustment of 0.5, and 200 µm² hole-fill cleanup with no size adjustment. Cell death (apoptosis and necrosis) was quantified from YOYO-1 green object counts using adaptive segmentation with a threshold adjustment (GCU) of 10, edge split enabled with edge sensitivity -45, 50 µm² hole-fill cleanup with one-pixel size adjustment, and filters for area (minimum 30 µm², maximum 6,000 µm²).

### IGF2 ELISA

For IGF2 quantification, cells were induced with doxycycline and transfected with siRNAs as described for the cell proliferation assays. Three days after transfection, conditioned medium was collected, and cells in each well were counted using a Countess™ automated cell counter (Thermo Fisher Scientific). Dead cells were removed by centrifugation, and medium aliquots were flash-frozen in liquid nitrogen. Frozen aliquots were thawed on ice, and IGF2 concentrations were measured using the Human IGF-II/IGF2 Quantikine ELISA Kit (R&D Systems) according to the manufacturer’s instructions. IGF2 concentrations were normalized to viable cell counts, and results were visualized using GraphPad Prism (v10.6.1, GraphPad Software).

### Histone PTM analysis by LC-MS/MS from cell lines

Cells were cultured on 10-cm dishes for 3 days prior to harvest. Approximately 5 × 10^6^ cells were washed twice with ice-cold PBS, collected in PBS supplemented with protease inhibitor cocktail (Sigma-Aldrich, P8340) and 10 mM sodium butyrate (Sigma-Aldrich, 303410), and flash frozen in liquid nitrogen. Histones were subsequently extracted and prepared for chemical derivatization and digestion as described previously^40,41^. In brief, lysine residues were propionylated using a reagent mixture of acetonitrile (Sigma-Aldrich, 34998) and propionic anhydride (Sigma-Aldrich, 8006080100) (3:1, v/v) at a 1:2 reagent-to-sample ratio. The reaction pH was adjusted to 8.0 with ammonium hydroxide (Sigma-Aldrich, 221228). The propionylation was repeated twice to ensure complete derivatization and samples were dried under vacuum. The derivatized histones were then digested overnight with 0.5 ug trypsin (Promega, V5113) (1:50, w/w) in 50 mM ammonium bicarbonate buffer at 37 °C. The resulting peptides were subjected to N-terminal propionylation (performed twice), dried again, and desalted using self-packed C18 stage tips. Purified peptides were reconstituted in 0.1% formic acid prior to LC-MS/MS analysis.

Peptide separation was performed on a Vanquish Neo UHPLC system coupled to an Orbitrap Exploris 240 mass spectrometer (Thermo Scientific). Samples were maintained at 7 °C and loaded onto an Easy-Spray™ PepMap™ Neo C18 column (2 µm, 75 µm × 150 mm). Chromatographic separation was achieved using a linear gradient from 2 to 32% solvent B (0.1% formic acid in acetonitrile) in solvent A (0.1% formic acid in water) over 48 min, followed by 42 to 98% solvent B over 12 min, at 300 nL/min.

Mass spectrometry data were acquired in data-independent acquisition (DIA) mode. Each acquisition cycle included one full MS scan followed by 35 DIA MS/MS scans with 24 m/z isolation windows spanning 295–1100 m/z. Full MS scans were collected in the Orbitrap mass analyzer at 60,000 resolution across 290–1100 m/z in positive profile mode, with automatic maximum injection time and an AGC target of 300%. MS/MS spectra were acquired after HCD fragmentation at 30 normalized collision energy (NCE), 1000% AGC target, and 60 ms maximum injection time.

Raw data were processed and quantified using EpiProfile (version 2.0)^42,43^. PTM ratios, defined as the MS1 area under the curve (AUC) of each modified peptide normalized to the sum of the unmodified and all modified forms of that peptide, were imported into R (www.R-project.org) for statistical analysis and visualization. Differential enrichment of PTMs was determined using the limma package (version 3.60.6)^44^ after log2 transformation of PTM ratios. P-values were adjusted using the Benjamini-Hochberg method to control the false discovery rate. PTMs with an adjusted p-value < 0.05 and |log₂FC| > 0.5 were considered significant. Data visualization, including heatmaps and volcano plots, was performed using ggplot2^45^ in R.

### Histone PTM analysis by LC-MS/MS from tumor samples

Flash-frozen human PN and MPNST samples were obtained from the Johns Hopkins University NF1 Biospecimen Repository (courtesy of Dr. Christine Pratilas). Tumor tissues were washed twice in ice-cold PBS and lysed using the Quick-DNA Microprep kit (Zymo Research) according to the manufacturer’s instructions. Briefly, tissues were homogenized with a mixture of 2.0 mm and 0.1 mm ceramic beads using a FastPrep-24™ bead homogenizer (MP Biomedicals) at 6.5 m/s for 60 s cycles until complete homogenization. Lysates were then sonicated with a probe sonicator (Fisherbrand, Thermo Scientific) for 10 s at 25% amplitude. Non-lysed debris was removed by centrifugation (twice), and the resulting supernatant was processed following the kit protocol. Proteins were precipitated with ice-cold acetone overnight, washed with 95–100% ethanol, and resuspended in 5% SDS in 50 mM triethylammonium bicarbonate buffer (TEAB, Sigma-Aldrich). Protein concentration was determined by BCA assay (Thermo Scientific) and lysates were used for histone PTM or total proteome analysis.

For histone PTM analysis, histone derivatization and trypsin digestion were performed in-gel as previously described^28,46^, with minor modifications. Briefly, 100 µg of protein lysate per sample in Laemmli buffer (Bio-Rad, 610737) containing 2.5% β-mercaptoethanol was loaded into two lanes of a 16% Tris-glycine SDS-PAGE gel. Gels were stained with GelCode™ Blue Safe Protein Stain (Thermo Scientific, 24596) for 15 min and destained in deionized water for 1 h. Histone bands were excised, cut into ∼1 mm^3^ cubes, and washed once in Milli-Q water for 1 h, followed by two 30 min washes in 50% acetonitrile. A third wash was performed overnight in 50% acetonitrile with shaking until all visible dye was removed.

Gel pieces were dehydrated with 100% acetonitrile twice, then derivatized for 20 min by adding 100 mM ammonium bicarbonate followed by propionic anhydride (1:2, v/v). After washing twice with 100 mM ammonium bicarbonate and shrinking again in 100% acetonitrile, the derivatization step was repeated. Gel pieces were then rehydrated in 50 mM ammonium bicarbonate containing 12.5 ng/µL trypsin for 45 min on ice and digested overnight at room temperature. The next day, peptides were extracted with 50% acetonitrile and dried under vacuum. N-terminal propionylation and all subsequent processing steps were performed as described for the cell line samples.

### Total proteome analysis

Cells were cultured on 10-cm dishes for 3 days prior to harvest. Approximately 5 × 10^6^ cells were washed twice with ice-cold PBS, collected in PBS supplemented with protease inhibitor cocktail (Sigma-Aldrich, MO, P8340) and 10 mM sodium butyrate (Sigma-Aldrich, MO, 303410), and flash frozen in liquid nitrogen. Cells were lysed in 5% SDS in 50 mM triethylammonium bicarbonate buffer (TEAB, Sigma-Aldrich, MO) and sonicated with a probe sonicator (Fisherbrand, Thermo Scientific, MA) for 10 sec at 20% power. Protein concentration was measured using BCA assay (Thermo Scientific, MA), and 50-100 ug of protein from cell lines or tumors were used for further processing. Reduction and alkylation of proteins were performed by adding 5 mM final concentration of tris(2-carboxyethyl)phosphine hydrochloride (TCEP, Sigma-Aldrich, MO) at 55 °C for 15 min and 20 mM final concentration of iodoacetamide (IAA, Sigma-Aldrich, MO) at room temperature for 30 min in the dark. The proteins were further loaded to the S-trap micro columns (ProtiFi, NY) and cleaned based on the manufacturer protocols. The proteins were digested with 1 μg of trypsin (Promega, WI) and 0.2 ug of Lys-C (FUJIFILM Wako Chemicals USA, VA) at 37 °C overnight. The digested peptides were eluted using 50mM TEAB, 0.2% formic acid (FA, Fisher Scientific, NH), and 50% acetonitrile (ACN, Thermo Scientific, MA). Peptides were reconstituted in 0.1% FA and quantified using Pierce Colorimetric Peptide Assay (Thermo Scientific, MA).

500 ng of peptide was characterized using a nanoACQUITY ultrahigh-pressure liquid chromatography (UPLC) System (Waters, MA) coupled with the ZenoTOF 7600 mass spectrometer (SCIEX, MA). The analytes were separated on a Phenomenex Kinetex XB C18 column (2.6 μm, 0.3 x 150 mm, Phenomenex, CA) at a flow rate of 10 μL/min, and the column temperature was set at 45 °C. A solution of water containing 0.1% FA and acetonitrile containing 0.1% FA were used as solvents A and B, respectively. The chromatography gradient consisted of 2% solvent B over 0-1 min, 2-30% solvent B over 1-46 min, 30-80% solvent B over 46-47 min, 80% solvent B over 47-49 min, and finally equilibrated with 2% solvent B over 5 min. The ZenoTOF 7600 was equipped with an OptiFlow Turbo V ion source and operated in SWATH mode with Zeno trap activated. The source conditions were as follows: ionization voltage: 5000 V, positive polarity, temperature: 200 °C, ion source gas 1: 20 psi, ion source gas 2: 60 psi, curtain gas: 35 psi. The Zeno SWATH DIA method consisted of 85 variable-width SWATH DIA windows that spanned the mass range 399.5-903.5 m/z, and the acquisition settings were as follows: MS1 accumulation time: 100 ms, MS1 m/z range: 400 to 1500, MS2 accumulation time: 13 ms, MS2 m/z range: 140 to 1800. For the gas phase fractionation (GPF) library, pooled sample was analyzed five times using the same settings as above but with following MS1 m/z ranges: 395 to 505, 495 to 605, 595 to 705, 695 to 805, 795 to 910, MS2 accumulation time: 15 ms.

All raw data were processed with Spectronaut (v20.2)^47^ using the directDIA mode supplied with the GPF library. The peptides were searched using default parameters: Trypsin/P, two missed cleavages allowed, toggle N-terminal M, fixed modification: C carbamidomethylation, variable modification: N-terminal acetylation, M oxidation, peptide length range: 7 to 52 amino acids. The proteins were mapped against the Homo sapiens UniProt canonical sequence database (UP000005640). Peptide and protein group quantities were imported into R for statistical analysis and visualization. Differential enrichment of proteins was determined using MSstats (v4.12.1)^48^. Proteins with an adjusted p-value < 0.05 and |log₂FC| > 0.5 were considered significant. Data visualization, including heatmaps and volcano plots, was performed using ggplot2 in R.

### ChIP-seq sample preparation

Cells were cultured on 15-cm dishes for 3 days prior to harvest in two biological replicates. Approximately 1 × 10^7^ cells were crosslinked with 1% (v/v) formaldehyde for 10 min at room temperature. Crosslinking was quenched by adding glycine to a final concentration of 120 mM and incubating for 10 min at room temperature. Cells were washed twice with ice-cold PBS, scraped from the plates, pelleted, washed twice again in PBS, and flash-frozen in liquid nitrogen. For ChIP-seq of Dox-inducible cells, TRE3G-SUZ12-90-8TL cells were cultured in standard growth medium supplemented with 1 µg/mL puromycin and 100 ng/mL doxycycline for 4 days. The medium was then replenished, and cells were cultured for an additional 3 days. After a total of 7 days of doxycycline induction, approximately 1 × 10^7^ cells were crosslinked, harvested, and processed as described for other cell lines.

Thawed pellets were resuspended in 500 µL cell lysis buffer (5 mM PIPES-pH 8.5, 85 mM KCl, 1% (v/v) IGEPAL CA-630, 50 mM NaF, 1 mM PMSF, 1 mM Phenylarsine Oxide, 5 mM Sodium Orthovanadate, EDTA-free Protease Inhibitor tablet) and incubated 30 minutes on ice. Samples were centrifugated and pellets resuspended in 500ul of nuclei lysis buffer (50 mM Tris-HCl pH 8.0, 10 mM EDTA, 1% (w/v) SDS, 50 mM NaF, 1 mM PMSF, 1 mM Phenylarsine Oxide, 5 mM Sodium Orthovanadate and EDTA-free protease inhibitor tablet) and incubated 30 minutes on ice.

Chromatin was sonicated on a BioRuptor UCD-300 (Diagenode) at maximum intensity for 60 cycles (10 s on, 20 s off), with centrifugation every 15 cycles and cooling by 4°C water circulation. Sonication efficiency was confirmed by agarose gel electrophoresis of a de-crosslinked and purified aliquot, targeting a fragment size of 150–500 bp. After sonication, chromatin was diluted to reduce the SDS concentration to 0.1% and concentrated using Nanosep 10K OMEGA filters (Cytiva). To enable normalization across samples, 2% of sonicated *Drosophila melanogaster* S2 cell chromatin was spiked in prior to immunoprecipitation.

ChIP reactions for histone modifications were performed using the Diagenode SX-8G IP-Star Compact system. Dynabeads Protein A (Invitrogen) were pre-washed and incubated with specific antibodies and 1.5 million cell equivalents of sonicated chromatin in the presence of protease inhibitors for 10 h. Antibodies used were rabbit monoclonal anti-H3K27me3 (Cell Signaling Technology, #9733), rabbit monoclonal anti-H3K36me2 (Cell Signaling Technology, #2901), rabbit polyclonal anti-H3K27ac (Active Motif, #39133), and rabbit polyclonal anti-H4K16ac (Active Motif, #39167). Immunocomplexes were washed for 20 min using the wash buffers provided in the Diagenode iDeal ChIP-seq kit for Histones.

Crosslinks were reversed by adding 4 µL 5 M NaCl and incubating at 65°C for 4 h. Samples were then treated with 2 µL RNase Cocktail at 65°C for 30 min followed by 2 µL Proteinase K at 65°C for 30 min. DNA was purified using the QIAGEN MinElute PCR Purification Kit according to the manufacturer’s protocol. Input DNA (equivalent to ∼50,000 cells) was processed in parallel following the same de-crosslinking and purification procedure.

ChIP-seq libraries were prepared using the KAPA HyperPrep Kit (Roche, #07962363001) with 8-bp dual-index adapters (Illumina TruSeq UDI v1, IDT for Illumina) following the manufacturer’s instructions. Libraries were sequenced on an Illumina NovaSeq X platform (paired-end 150 bp) at the McGill Genome Centre (Montreal, Canada).

### ChIP-seq data-analysis

Raw ChIP-seq reads were processed using a standardized and reproducible workflow following ENCODE and modENCODE best-practice guidelines. Adapter trimming and quality control were performed with fastp (v0.23.4)^49^, enabling removal of low-quality bases, adapter sequences, and overrepresented reads. Read quality metrics were assessed using FastQC (v0.12.1) prior to and following trimming.

For alignment, cleaned reads were mapped to the human reference genome (hg38) using BWA-MEM (v0.7.17)^50^ with default parameters. Alignment files were sorted and indexed using SAMtools (v1.20)^51^, and PCR duplicates were marked and removed with Picard (v3.1.1; Broad Institute). Mapping statistics were summarized using SAMtools flagstat and Qualimap (v2.3)^52^.

Spike-in (Drosophila melanogaster, dm6) normalization was incorporated to correct for global differences in ChIP efficiency across samples, following the ENCODE normalization framework. Spike-in reads were separated by reference genome and quantified independently before normalization of ChIP signal intensities.

Filtered and duplicate-marked BAM files were converted into normalized coverage tracks using deepTools (v3.5.5)^53^. The bamCoverage function was used to generate CPM-normalized bigWig files (bin size 50 bp) with read extension to 200 bp, and bigwigAverage function was used to generate average bigwig files from biological replicates. Genomic coverage and library complexity were further assessed using deepTools plotFingerprint and plotCoverage modules.

Peaks were identified using MACS2 (v2.2.9.1)^54^ in both narrow-peak and broad-peak modes with -- nomodel and --nolambda parameters to account for variable fragment lengths and background bias. Genome size parameters were set to mm for mouse or hs for human samples. Broad peaks were identified using a false discovery rate (FDR) threshold of 0.1, while narrow peaks were reported at default thresholds. Peak files were converted to bigBed format for genome browser visualization using UCSC bedToBigBed.

QC and library complexity metrics were evaluated using deepTools, RSeQC^55^, and read_distribution.py, including coverage uniformity, duplication rates, and gene body coverage bias. Peak reproducibility was verified through cross-correlation metrics (phantompeakqualtools) and replicate concordance following ENCODE IDR guidelines^56^.

For H3K27me3 and H3K27ac, broad peaks that were identified in both biological replicates were considered representative for the given condition and used in the analysis. Numbers of overlapping peaks in different conditions were determined using bedtools (v.2.31.0)^57^ *intersect* tool. DESeq2^58^ with Diffbind^59^ R package was used to determine differentially enriched peaks. Peaks with FDR <0.05 and |log2FC| >1 were considered as significantly differentially enriched. Peaks were annotated using *annotatePeaks* tool in HOMER^60^. Differential H3K36me2 ChIP-seq signal from gene bodies was analyzed using *analyzeRepeats* tools in HOMER and edgeR^61^. Genes with with adjusted p-value <0.05 and |log2FC| >1 were considered as significantly differentially enriched. Gene sets with changed histone PTMs were subjected to pathway analysis with DAVID functional annotation tool^62,63^ for direct biological process GO-terms.

For data visualization, heatmaps and average profiles were generated using deepTools *computeMatrix*, *plotHeatmap* and *plotProfile* tools. Scatter plots were generated using ggplot2^45^ in R. Chromatin tracks were visualized in Integrative Genomics Viewer genome browser (IGV, Broad Institute) and assembled in Photoshop and Illustrator (Adobe). Column graphs for genomic distributions and GO-terms were generated with GraphPad Prism (v10.6.1, GraphPad Software).

### RNA-seq

Cells were cultured on 6-well plates for 3 days prior to harvest in three biological replicates. Total RNA from approximately 1 × 10^6^ cells per sample was extracted with RNeasy Plus Mini kit (Qiagen) following manufacturer’s instructions. For Dox-inducible cells, TRE3G-SUZ12-90-8TL cells were cultured in standard growth medium supplemented with 1 µg/mL puromycin and 100 ng/mL doxycycline for 4 days in three biological replicates. The medium was then replenished, and cells were cultured for an additional 3 days. After a total of 7 days of doxycycline induction, RNA from approximately 1 × 10^6^ cells per sample was extracted and processed as for the other cell lines. Library preparation and raw data processing were performed at the Genome Technology Access Center (GTAC), Washington University in St. Louis.

Samples were prepared according to library kit manufacturer’s protocol, indexed, pooled, and sequenced on an Illumina NovaSeq X Plus. Basecalls and demultiplexing were performed with Illumina’s DRAGEN and BCLconvert version 4.2.4 software. RNA-seq reads were then aligned to the Ensembl_R113 primary assembly with STAR version 2.7.11b^64^. Gene counts were derived from the number of uniquely aligned unambiguous reads by Subread:featureCount version 2.0.8^65^. Isoform expression of known Ensembl transcripts were quantified with Salmon version 1.10.0^66^. Sequencing performance was assessed for the total number of aligned reads, total number of uniquely aligned reads, and features detected. The ribosomal fraction, known junction saturation, and read distribution over known gene models were quantified with RSeQC version 5.0.4^55^.

All gene counts were then imported into the R/Bioconductor v4.4.0 package EdgeR^61^ and TMM normalization size factors were calculated to adjust for samples for differences in library size.

Ribosomal genes and genes not expressed in at least 3 samples greater than 1 count-per-million were excluded from further analysis. The TMM size factors and the matrix of counts were then imported into the R/Bioconductor package Limma^44^. Weighted likelihoods based on the observed mean-variance relationship of every gene and sample were then calculated for all samples with the voomWithQualityWeights^67^ function and were fitted using a Limma generalized linear model.

Genes with with adjusted p-value <0.05 and |log2FC| >1 were considered as differentially enriched. Differentially enriched gene sets were subjected to pathway analysis with DAVID functional annotation tool^62,63^ for direct biological process GO-terms. Column graphs for GO-terms were generated with GraphPad Prism (v10.6.1, GraphPad Software). Average bigwig files from biological replicates were generated using deepTools (v3.5.3)^53^ bigwigAverage function. Chromatin tracks were visualized in Integrative Genomics Viewer genome browser (IGV, Broad Institute) and assembled in Photoshop and Illustrator (Adobe). Scatter plots were generated using ggplot2^45^ in R.

### Public dataset analysis

Raw fastq files for primary MPNST transcriptomes were retrieved from the Gene Expression Omnibus (GSE206527)^31^ and the European Genome-phenome Archive (EGAD00001008608). Access to the EGA dataset generated by the Genomics of MPNST (GeM) consortium^18^ was granted through a Data Access Agreement with the Neurofibromatosis Research Initiative. In total, 166 MPNST transcriptomes were analyzed. Raw fastq files with disordered read-pairs were repaired using the *repair.sh* script from the BBTools toolkit^68^. After Fastq repair, quality control analysis, and exclusion of MPNST samples with unknown PRC2 status, 150 MPNST transcriptomes were retained for further analysis.

Adapter sequences were trimmed from fastq files using cutadapt^69^. Paired reads were aligned to the *hg38* reference genome using the STAR aligner^64^ (two-pass mode) with the Gencodev26^70^ annotation. RNA strandedness was assessed using *infer_experiment.py* (RSeQC)^55^, and gene-level counts were generated using *featureCounts* (Subread)^65^ using the same annotation.

Gene-level counts were normalized by library size using *DESeq2*^58^. Counts per million (CPM) values were calculated using *edgeR*^61^, and genes with CPM < 2 in fewer than 10 samples were excluded prior to normalization. Samples with a mean Pearson correlation < 0.2 relative to the cohort were excluded from further analysis. Normalized counts were subjected to a variance stabilizing transformation (VST) before downstream visualization and analyses.

## Data availability

Genomics and proteomics data will be made publicly available after manuscript peer review.

## Acknowledgements

We thank the GTAC and GESC cores at Washington University School of Medicine for RNA-seq, mycoplasma testing and STR profiling services. We thank the Research Infrastructure Services at Washington University in St. Louis for providing computational infrastructure.

