## Supplementary Figures for "Integrated Proteomic and Epigenomic Analysis Reveals IGF2 as a Vulnerability in PRC2-Deficient Malignant Peripheral Nerve Sheath Tumors"

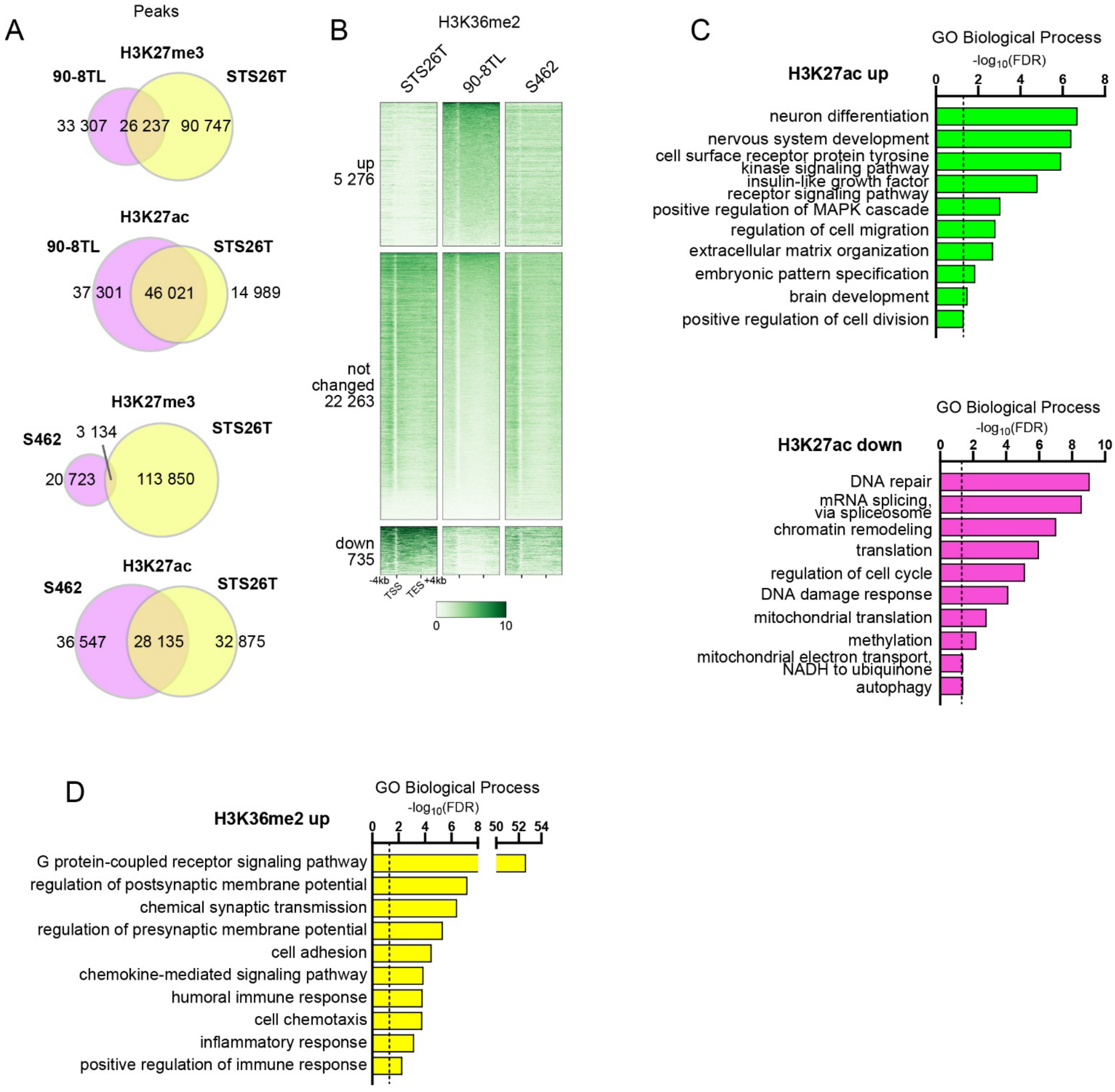


**Supplementary Figure S1. Related to main Figure 1. (A)** Overlap of H3K27me3 and H3K27ac ChIP-seq peaks in 90-8TL, STS26T and S462 cells. **(B)** H3K36me2 signal at gene bodies clustered as upregulated (adj. $P < 0.05$, $\log_2FC > 1$), downregulated (adj. $P < 0.05$, $\log_2FC < -1$), or unchanged in PRC2-deficient 90-8TL cells relative to PRC2-intact STS26T cells. The same gene regions are shown for PRC2-deficient S462 cells. TSS; transcription start site, TES; transcription end site. **(C)** Significantly enriched Gene Ontology (GO) Biological Processes (FDR < 0.05) for genes with up- or downregulated H3K27ac. **(D)** GO Biological Processes (FDR $< 0.05$) for genes with upregulated H3K36me2.

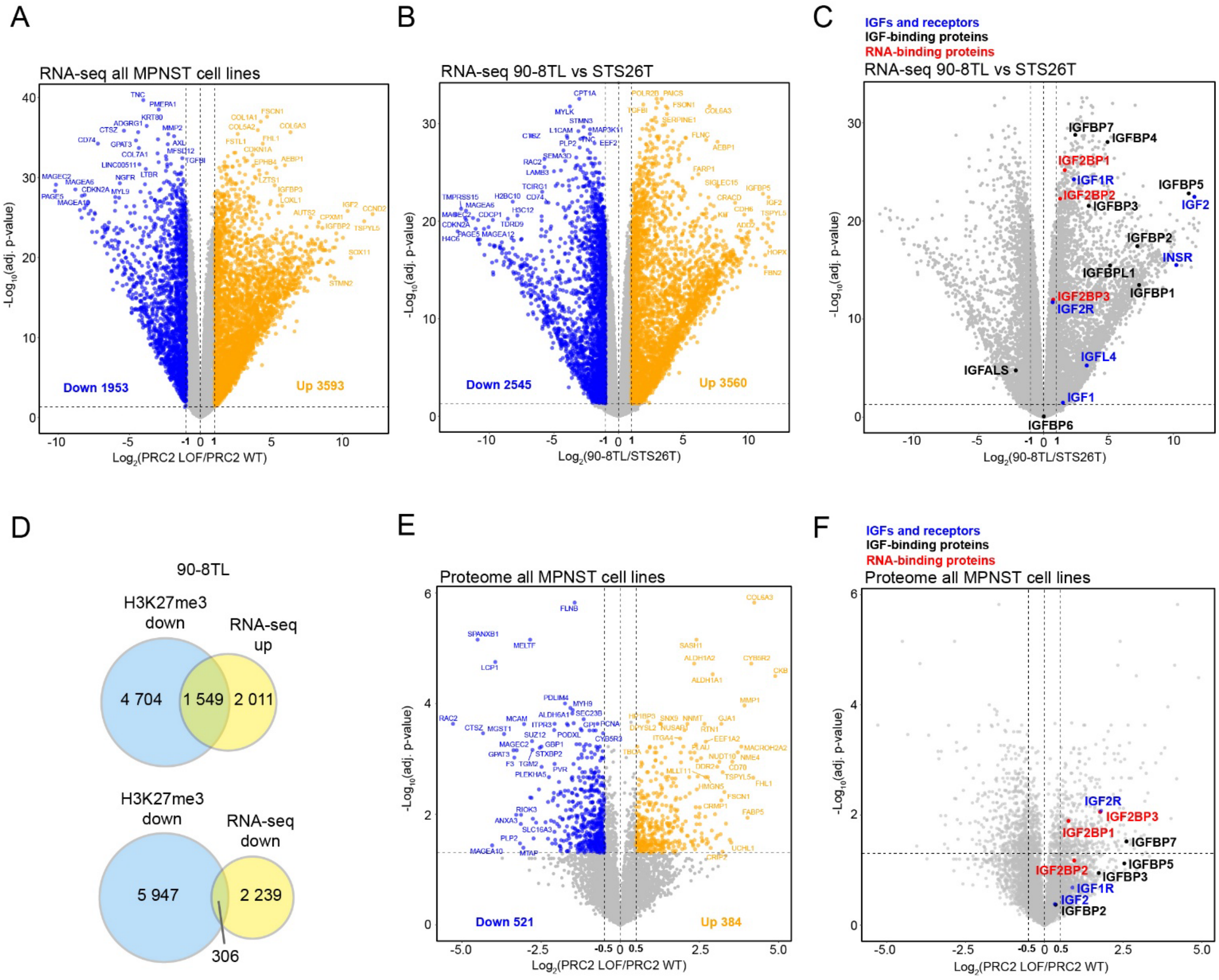


**Supplementary Figure S2. Related to main Figure 2. (A)** Significantly altered genes from the RNA-seq comparison between all PRC2-deficient and PRC2-intact cell lines. **(B)** Significantly altered genes from the RNA-seq comparison between 90-8TL versus STS26T cells. **(C)** Volcano plot highlighting IGF2-signaling -related genes from the RNA-seq comparison in B. **(D)** Overlap of genes with downregulated H3K27me3 and up- or downregulated mRNA expression in 90-8TL cells compared to STS26T. **(E)** Significantly altered proteins from the quantitative proteomics comparison between all PRC2-deficient and PRC2-intact cell lines. **(F)** Volcano plot highlighting IGF2-signaling -related proteins from the proteomics comparison in E.

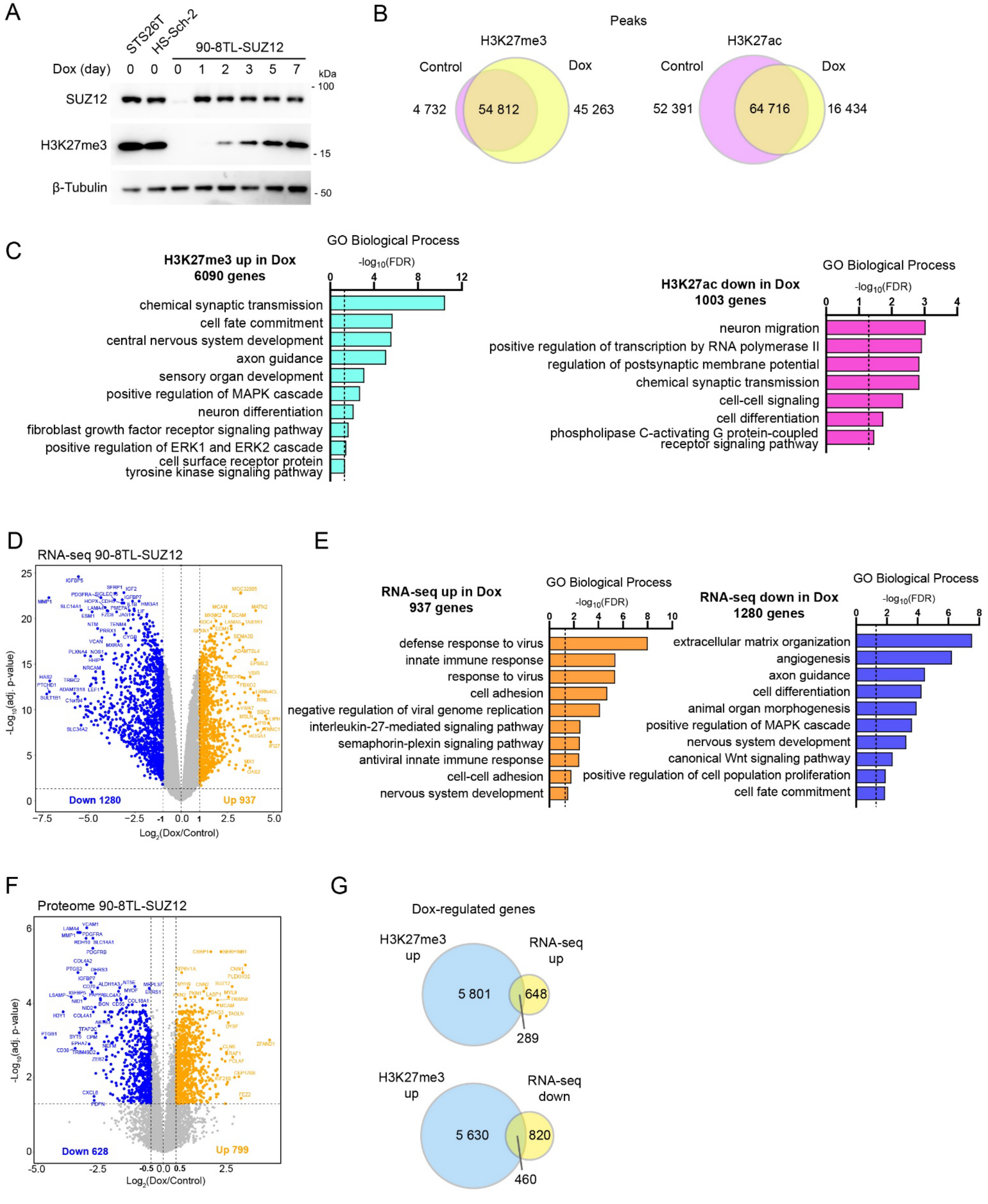


**Supplementary Figure S3. Related to main Figure 3. (A)** Western blot showing the level of SUZ12 and H3K27me3 in PRC2-intact cells (STS26T and HS-Sch-2) and in Doxycycline (Dox) -inducible SUZ12-restored 90-8TL cells during 0-7 days of Dox induction. **(B)** Overlap of H3K27me3 and

H3K27ac ChIP-seq peaks in control and Dox-treated cells. **(C)** Significantly enriched Gene Ontology (GO) Biological Processes (FDR < 0.05) for genes with upregulated H3K27me3 or downregulated H3K27ac in Dox-treatment. (**D**) Significantly altered genes from the RNA-seq comparison between Dox and control-treated cells. (**E**) Significantly enriched GO Biological Processes (FDR < 0.05) for up- and downregulated genes in D. **(F)** Significantly altered proteins from the quantitative proteomics comparison between Dox and control-treated cells. **(G)** Overlap of genes with upregulated H3K27me3 and up- or downregulated mRNA expression in Dox-treated cells.

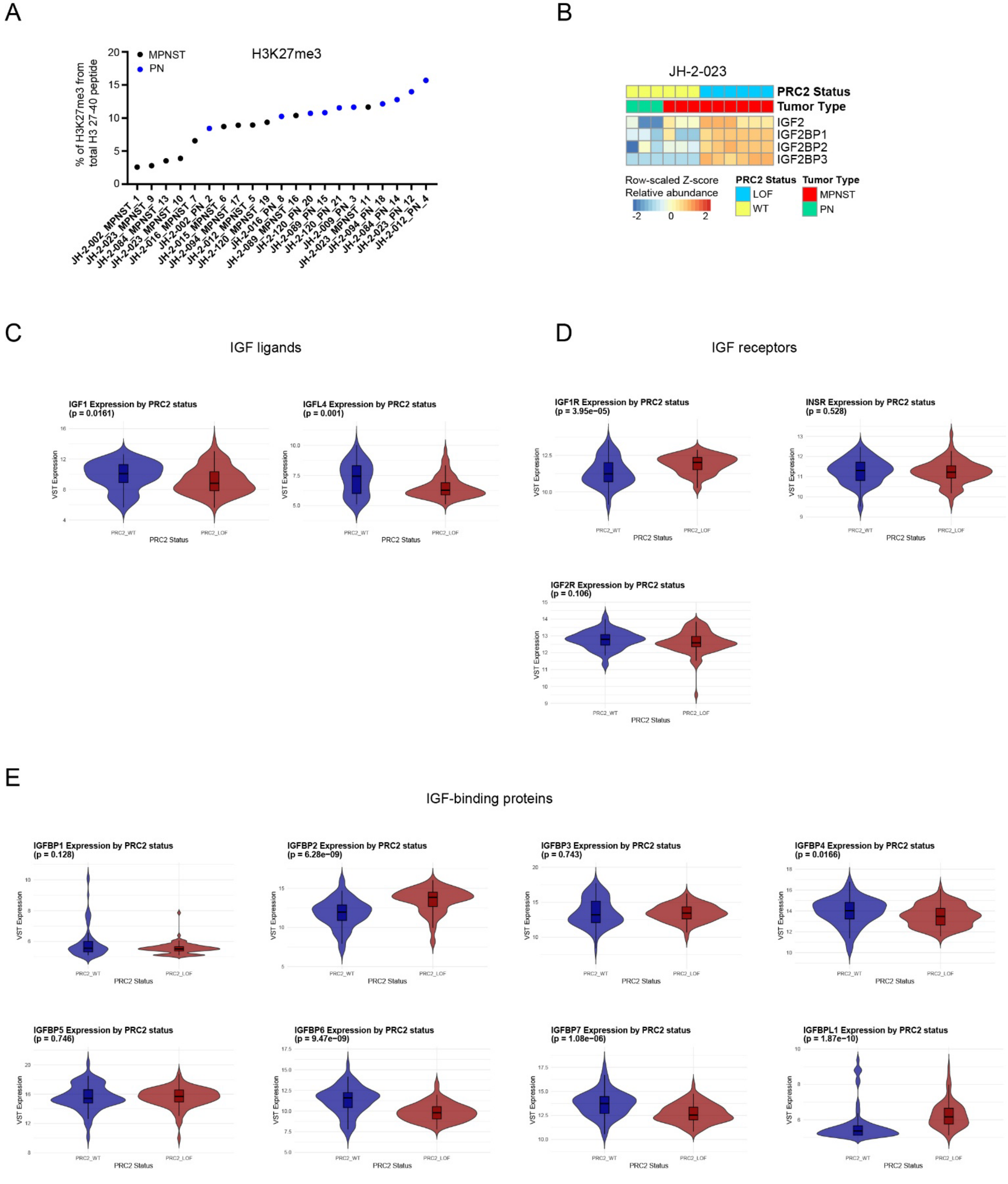


**Supplementary Figure S4. Related to main Figure 5. (A)** Relative H3K27me3 levels in human MPNSTs and PNs detected by global histone PTM MS analysis. **(B)** Protein expression of IGF2,

IGF2BP1, IGF2BP2 and IGF2BP3 in MPNST and PN tumors from the JH-2-023 patient measured by quantitative proteomics. Data represent three technical replicates. **(C)** Violin plots showing average mRNA expression of *IGF1* and *IGF4L* in PRC2-WT and -LOF tumors from the MPNST cohorts (GeM and GSE206527). Statistical significance was assessed by Wilcoxon test. **(D)** Same as (C) but for *IGF1R*, *INSR* and *IGF2R* genes. **(E)** Same as (C) and (D) but for IGFBP genes.
